# Identification of *srh-30* as a 2-nonanone receptor in *C. elegans*

**DOI:** 10.64898/2026.06.15.732464

**Authors:** Yan Jin, Alessandro Groaz, Heenam Park, Paul W. Sternberg

**Author notes:** Y.J. and A.G. contributed equally to this work.

## Abstract

*C. elegans* detects food, mates, and danger through a large repertoire of chemosensory receptors. The volatile odorant 2-nonanone is a well-established aversive stimulus that reliably elicits avoidance chemotaxis. However, the receptor(s) responsible for its detection have not been identified. Here, we leveraged CeNGEN single-cell expression profiles to identify candidate receptors required for sensing 2-nonanone. We narrowed the list of candidate chemoreceptor genes by selecting genes expressed in the aversive amphid neurons AWB and ASH while excluding genes expressed in other amphid sensory neurons. We tested available mutants from the *C. elegans* Genetics Center (CGC) and identified *srh-30* as a candidate required for normal 2-nonanone avoidance. Two independent *srh-30* null alleles generated using CRISPR-Cas9 confirmed that loss of *srh-30* causes a partial defect in 2-nonanone sensing. *srh-30* promoter-driven fluorescent reporter, mKate, localized to the distal tips of AWB sensory cilia, consistent with a direct sensory role. Finally, ectopic expression of *srh-30* in the attractive AWA neuron shifted the behavioral response to 2-nonanone toward neutrality, supporting *srh-30* as a sensory receptor. Together, these results identify *srh-30* as a sensory receptor mediating 2-nonanone-evoked avoidance and demonstrate a transcriptome-guided strategy for mapping odor receptors.

## Introduction

Animals rely on chemosensation to detect food, danger, mates, pathogens, and predators[1]. *Caenorhabditis elegans* provides a powerful and highly tractable system for studying how environmental cues are detected and transformed into behavior[2, 3]. Despite having only 302 neurons[4], *C. elegans* can discriminate a wide range of soluble and volatile chemicals and generate robust behavioral responses, including attraction, avoidance, and experience-dependent modulation of chemotaxis[5-8]. Its compact and fully mapped nervous system, stereotyped sensory neuron identities, and powerful genetic toolkit make it especially well-suited for linking specific odorants to their receptors, sensory neurons, neural circuits, and behavioral outputs[9, 10].

*C. elegans* detects many chemical cues through 12 bilateral pairs of amphid sensory neurons, which are functionally specialized[11]. AWA and AWC mediate attraction to many volatile odorants[7], whereas AWB and ASH contribute to avoidance of aversive volatile and noxious cues[3]. The molecular basis of olfaction in *C. elegans* is encoded in an unusually large repertoire of approximately 1,300 predicted seven-transmembrane chemoreceptor genes, distributed across many families, including the *srh, str, sra*, and *srg* classes, and expressed in partially overlapping, neuron-specific combinations[12-14]. Despite the depth of behavioral and anatomical characterization of *C. elegans* olfaction, only a small number of these receptors have been matched to specific ligands. The diacetyl receptor ODR-10, expressed in AWA, remains the textbook example of a deorphanized *C. elegans* odorant receptor[15], and the cognate receptors for most individual odorants, including most aversive volatiles, remain unknown[16-18].

The volatile ketone 2-nonanone, a microbial volatile produced by several bacterial species, is a well-established aversive odorant in *C. elegans*[19, 20]. However, the receptor or receptors responsible for its detection remain unknown[2]. Defining a 2-nonanone receptor would provide an entry point for dissecting how this odorant is detected and how receptor activation is coupled to avoidance behavior.

Prior laser-ablation, genetic, and calcium-imaging studies have implicated AWB, and to a lesser extent ASH, in 2-nonanone avoidance[19, 21-23]. However, identifying the relevant receptor from more than 1,300 predicted seven-transmembrane chemoreceptor candidates is experimentally challenging. Single-cell transcriptomic resources, particularly the CeNGEN atlas, now make it possible to prioritize candidates based on neuron-specific expression[24]. Here, we applied this transcriptome-guided strategy to search for a 2-nonanone receptor. Using CeNGEN expression profiles, we identified a short list of candidate chemoreceptors expressed in AWB and ASH while excluding receptors expressed in other amphid sensory neurons.

We then screened the corresponding mutant strains available from the *Caenorhabditis* Genetics Center for defects in 2-nonanone avoidance. This screen identified *srh-30* as a gene required for normal 2-nonanone avoidance. To confirm this result, we generated two independent CRISPR-Cas9 null alleles[25-27] of *srh-30* and found that both caused a partial but reproducible defect in 2-nonanone avoidance, while avoidance of 1-octanol was unaffected. Fluorescent reporters showed that *srh-30* is expressed in AWB and that SRH-30 protein localizes to the distal tips of AWB sensory cilia, consistent with a direct sensory role. Finally, ectopic expression of *srh-30* in the attractive AWA neuron, under the *odr-10* promoter, shifted the *srh-30* mutant response to 2-nonanone toward neutrality without affecting AWA-mediated attraction to diacetyl. Together, these results identify SRH-30 as a chemoreceptor contributing to 2-nonanone-evoked avoidance and illustrate a transcriptome-guided pipeline for matching orphan chemoreceptors to their cognate odorants.

## Results

### Transcriptome-guided candidate selection identifies *srh-30* as a candidate receptor for 2-nonanone avoidance

To identify candidate receptors for aversive volatile odorants, we first used CeNGEN single-cell expression profiles to prioritize chemoreceptor genes expressed in the aversive amphid neurons AWB and/or ASH. Because 2-nonanone avoidance has been linked primarily to AWB, with an additional contribution from ASH, we focused on receptors enriched in these neurons and excluded candidates with broader expression across other amphid sensory neurons. This filtering strategy reduced the candidate list from the genome-wide chemoreceptor repertoire to 16 genes, 7 of which had mutant strains available from the Caenorhabditis Genetics Center (CGC) (Fig. 1A, Supplementary Table 1).

**Figure 1.**
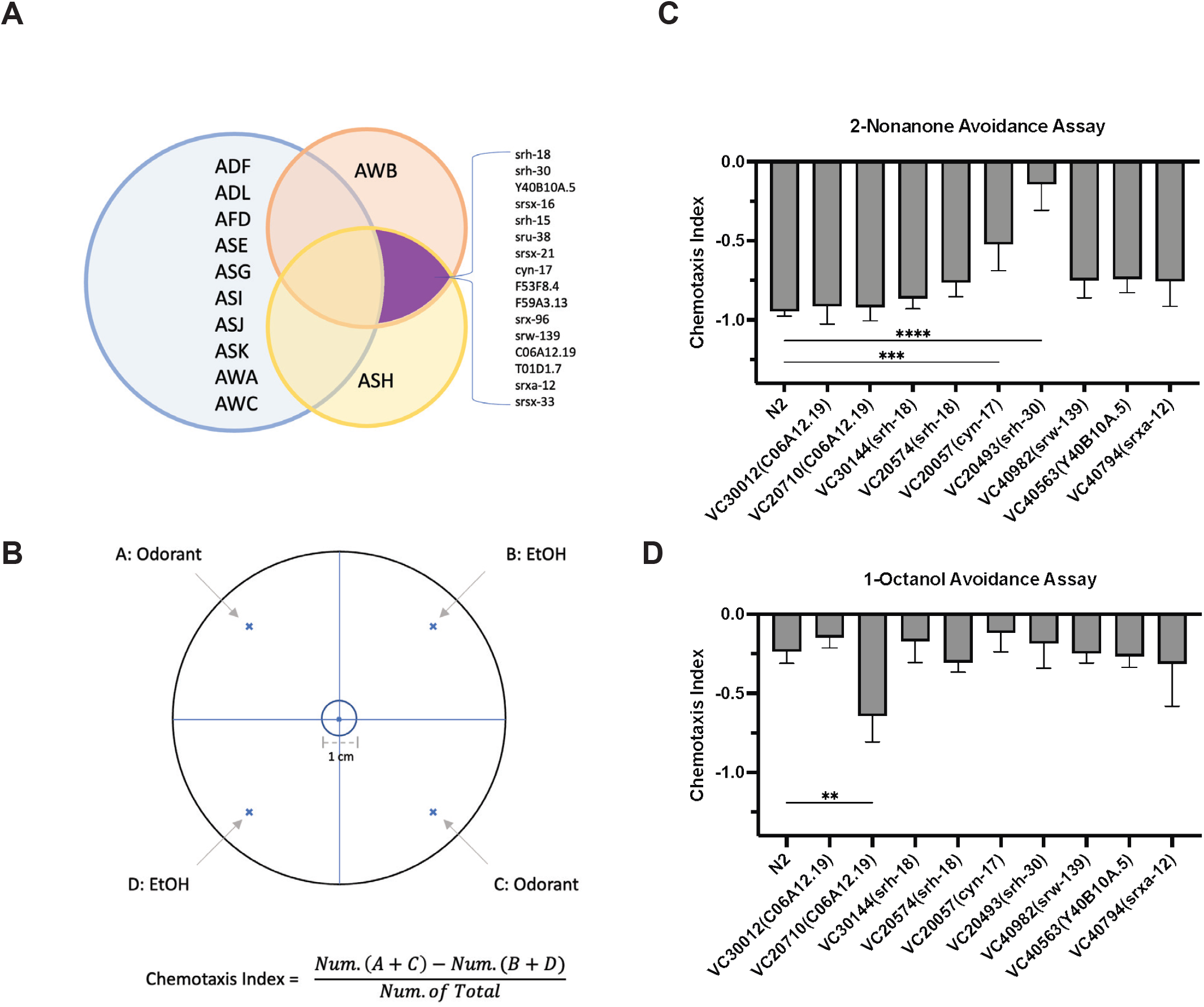
Identifying candidate chemoreceptors that required for sensing aversive odors using CeNGEN single cell RNA sequencing data. A) Venn diagram illustrating aversive odor receptor candidates that express in the aversive amphid neurons AWB and ASH, with minimal or no expression in other amphid sensory neurons using CeNGEN expression data. B) Schematic of the quadrant avoidance assay. A 10 cm Petri dish was divided into four quadrants (A to D). Odorant (2-nonanone or 1-octanol) was placed at positions A and C, and ethanol (EtOH) control was placed at positions B and D. Worms were placed at the center, and the chemotaxis index (CI) was calculated as CI = (Num.(A + C) ™ Num.(B + D)) / total animals scored. Animals remaining within the central circle (1 cm diameter) were excluded from scoring. C) Avoidance (chemotaxis) index to 2-nonanone for N2 and available mutant strains obtained from the CGC (strain IDs shown with the corresponding gene). VC20493 (*srh-30*) and VC20057 (*cyn-17*) show reduced avoidance to 2-nonanone. D) Avoidance (chemotaxis) index to 1-octanol for the same set of strains. VC20710 (*C06A12*.*19*) shows reduced avoidance to 1-octanol. Bars show mean ± SEM. Asterisks indicate statistical significance relative to N2 by one-way ANOVA with Dunnett’s multiple-comparisons test (* P < 0.05, ** P < 0.01, *** P < 0.001, **** P < 0.0001), ns indicates not significant.

We then screened the available mutant strains using a quadrant avoidance assay [28, 29], in which 2-nonanone or 1-octanol was placed in two opposing quadrants and ethanol controls were placed in the remaining quadrants (Fig. 1B). In this assay, animals that avoid the odorant accumulate preferentially in the ethanol quadrants, producing a negative chemotaxis index. Most candidate mutants retained robust avoidance to 2-nonanone. However, the *srh-30* mutant strain VC20493 showed significantly reduced 2-nonanone avoidance compared with wild-type N2 animals (Fig. 1C). The *cyn-17* mutant strain VC20057 also showed reduced avoidance in this primary screen, whereas the other tested candidate mutants did not produce comparable defects.

To determine whether these phenotypes reflected a specific defect in 2-nonanone chemotaxis rather than a general impairment in aversive odor responses, we tested the same set of strains with 1-octanol. In contrast to its reduced response to 2-nonanone, the *srh-30* mutant did not show a comparable defect in 1-octanol avoidance in the quadrant assay (Fig. 1D). Instead, the strongest 1-octanol phenotype was observed in the *C06A12*.*19* mutant strain VC20710.

Because *cyn-17* is annotated as a cyclophilin [30, 31] rather than a seven-transmembrane chemoreceptor, and because *C06A12*.*19* is not classified among the established *C. elegans* seven-transmembrane chemoreceptor families [12-14], we did not prioritize these genes as likely odorant receptor candidates. Together, these results suggest that loss of *srh-30* impairs behavioral avoidance of 2-nonanone without causing a broad defect in aversive chemotaxis. We therefore focused subsequent experiments on *srh-30*.

### Independent CRISPR null alleles confirm that *srh-30* is required for normal 2-nonanone avoidance

Because the initial CGC strain was derived from a chemically mutagenized background and may contain additional mutations [32], we next tested whether the 2-nonanone avoidance defect was specifically caused by loss of *srh-30*.

To do this, we generated two independent *srh-30* null alleles, *srh-30(sy1924)* and *srh-30(sy1925)*, using CRISPR-Cas9 genome editing (Fig. 2A) [27]. We then examined these mutants using a two-choice aversive chemotaxis assay, in which 2-nonanone was placed on one side of the plate and ethanol control was placed on the opposite side (Fig. 2B). Both independent *srh-30* null mutants showed significantly reduced avoidance of 2-nonanone compared with wild-type N2 animals (Fig. 2C). These results confirmed that *srh-30* is required for normal behavioral responses to 2-nonanone. However, the defect was partial rather than complete, suggesting that additional receptors or parallel sensory pathways may also contribute to 2-nonanone avoidance.

**Figure 2.**
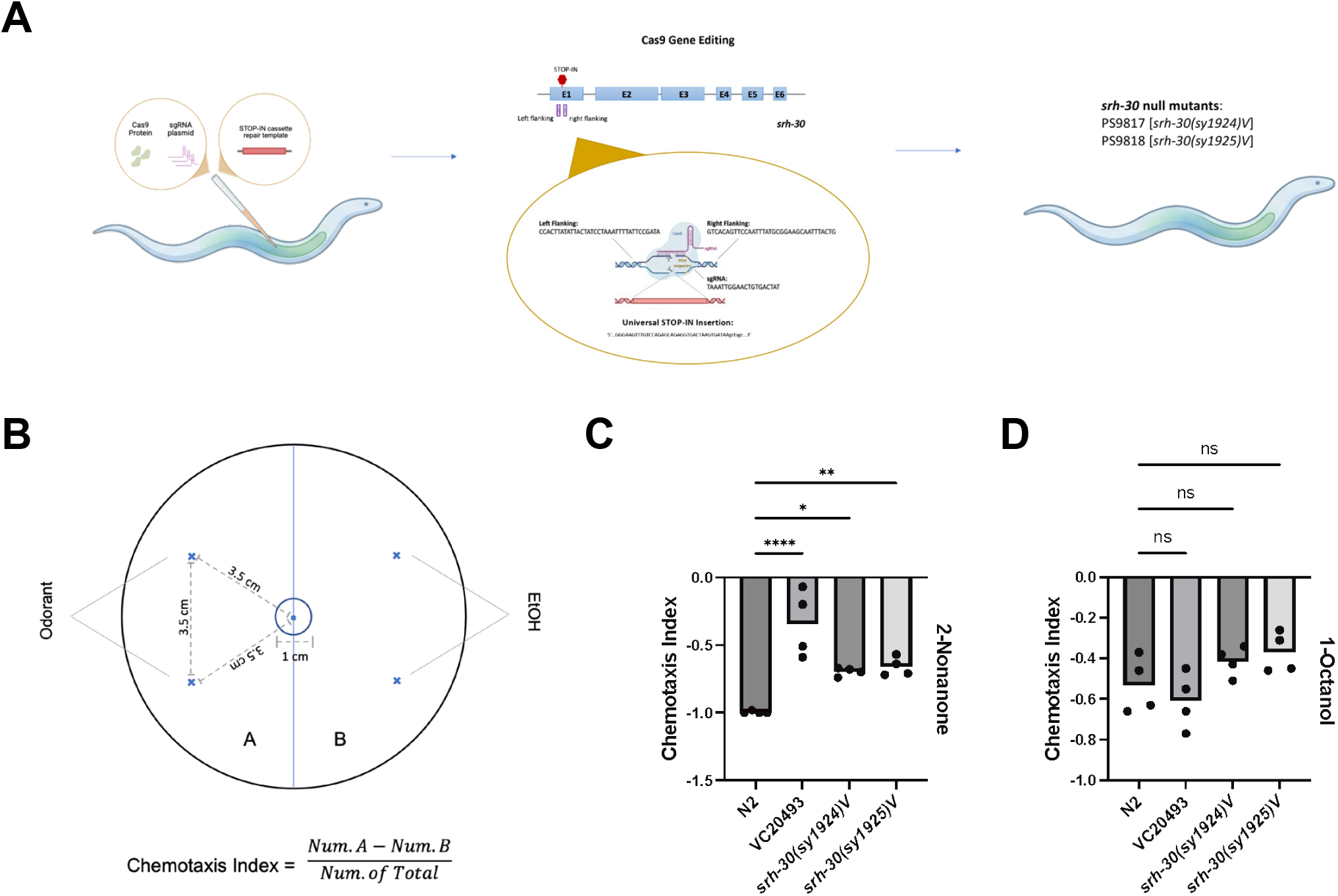
*srh-30* null mutants are defective in Chemotaxis to 2-nonanone. A) Schematic of CRISPR Cas9 gene editing used to generate two independent *srh-30* null alleles, resulting in *srh-30(sy1924)V [PS9817]* and *srh-30(sy1925)V [PS9818]*. B) Schematic of the two-choice aversive chemotaxis assay. A 10 cm Petri dish was divided into two halves (A and B). One microliter of odorant (2-nonanone or 1-octanol) was placed on the odorant side, and 1 µL ethanol (EtOH) was placed on the control side. Worms were placed at the center, and the chemotaxis index (CI) was calculated as CI = (Num. worms in A ™ Num. worms in B) / total worms scored. Animals remaining within the central circle (1 cm diameter) were excluded from scoring. C) Chemotaxis index to 2-nonanone. *srh-30(sy1924)V [PS9817]* and *srh-30(sy1925)V [PS9818]* show significantly reduced avoidance compared with wild type *N2*. D) Chemotaxis index to 1-octanol. *srh-30(sy1924)V [PS9817]* and *srh-30(sy1925)V [PS9818]* do not differ significantly from wild type *N2*. Bars show mean ± SEM, dots indicate individual trials. Asterisks indicate statistical significance relative to *N2* (* P < 0.05, ** P < 0.01, *** P < 0.001, **** P < 0.0001), ns indicates not significant.

To determine whether the reduced 2-nonanone response reflected a general defect in aversive chemotaxis, we tested the same mutants with 1-octanol. Both *srh-30* null mutants avoided 1-octanol at levels not significantly different from wild-type animals (Fig. 2D). These results argue against a broad defect in aversive chemotaxis and indicate that *srh-30* is required for normal behavioral responses to 2-nonanone.

### *srh-30* is expressed in AWB neurons and SRH-30 localizes to the distal sensory cilia

*To* determine whether *srh-30* is expressed in the sensory neurons implicated in 2-nonanone avoidance, we generated a transcriptional reporter in which the *srh-30* promoter drives expression of mKate2 (Fig. 3A). Reporter expression was observed in a pair of amphid sensory neurons with morphology and position consistent with AWB. To confirm this identity, we crossed the *srh-30p::mKate2* reporter with an AWB marker line that uses the cGAL-UAS system previously developed in our lab[33, 34]. In this marker strain, the *str-1* promoter drives the cGAL transcriptional activator GAL4(sk)::VP64 in AWB neurons, which in turn activates a UAS::GFP effector. The strain also carries transgenic RFP markers, including *unc-122p::RFP* with the syIs666 driver and *ttx-3p::RFP* with the syIs300 effector, which were used to track transgenes rather than to define AWB identity. The mKate2 signal colocalized with the AWB GFP marker, indicating that *srh-30* is expressed in AWB neurons (Fig. 3B).

**Fig. 3.**
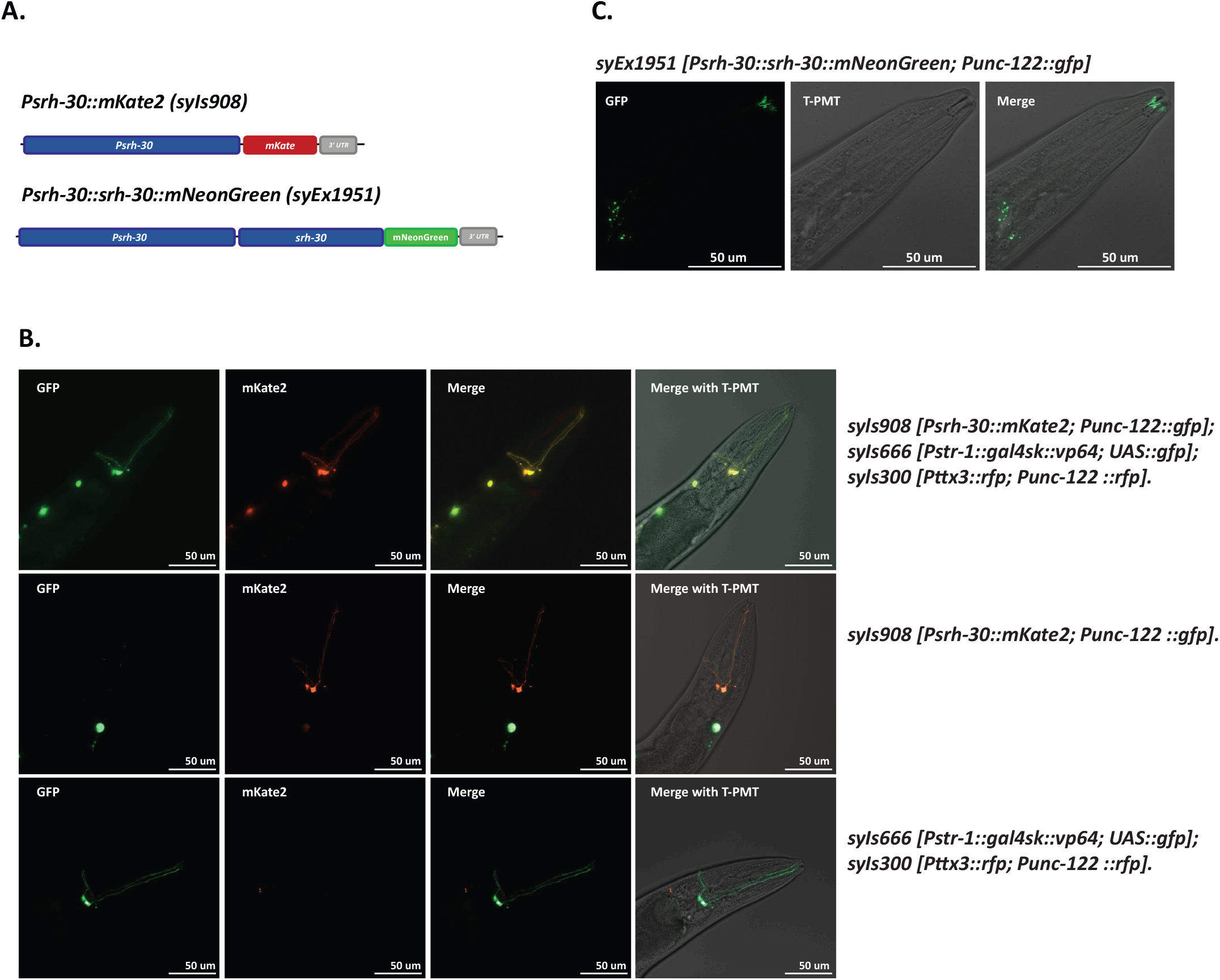
srh-30 Expresses in AWB Neurons and Localized at the Tips of Its Sensory Cilia. A) Schematic diagram of *srh-30* expression constructs. *srh-30* promoter (*Psrh-30*), 3Kb upstream of *srh-30* coding sequence, drives the expression of mKate2 *(syIs908). srh-30* promoter drives the expression of SRH-30 and mNeonGreen fusion protein *(syEx1951)*. B) SRH-30 expresses in AWB neurons. mKate2, which is driven by *srh-30* promoter (*syIs908*), colocalized with GFP expressed under the control of *str-1:cGAL*. C) SRH-30 is expressed at the tips of AWB cilia. mNeonGreen is fused to the C terminus of SRH-30 and signal was detected at the tips of AWB neurons (*syEx1951*).

Because chemosensory receptors are expected to function at sensory cilia, we next examined the subcellular localization of SRH-30. We generated a translational reporter in which SRH-30 was fused to mNeonGreen and expressed under the control of the *srh-30* promoter (Fig. 3A). SRH-30::mNeonGreen signal was enriched at the distal tips of AWB sensory cilia (Fig. 3C). This localization is consistent with a role for SRH-30 in odor detection and supports the model that SRH-30 functions as a receptor in AWB sensory cilia to mediate 2-nonanone avoidance.

### Ectopic expression of *srh-30* in AWA shifts behavioral responses toward neutrality

In *C. elegans*, amphid sensory neuron classes are functionally specialized, with different neuron classes detecting characteristic chemical cues and preferentially driving distinct behavioral outputs [2]. For example, AWB neurons mediate avoidance to aversive volatile cues, whereas AWA neurons mediate attraction to attractive odorants such as diacetyl. Previous receptor misexpression experiments showed that behavioral valence can depend on the sensory neuron in which an odorant receptor is expressed [15, 19]. Therefore, if SRH-30 functions as a receptor for 2-nonanone, expressing *srh-30* in the attractive AWA neuron could alter how the animal interprets 2-nonanone.

To test this idea, we expressed *srh-30* in AWA using the AWA-specific *odr-10* promoter in the *srh-30* null mutant background (Fig. 4A). Ectopic expression of *srh-30* in AWA shifted the behavioral response to 2-nonanone toward neutrality in both *srh-30(sy1924)* and *srh-30(sy1925)* mutant animals (Fig. 4B). Although this manipulation did not convert 2-nonanone into a strongly attractive cue, it significantly reduced avoidance, consistent with the idea that SRH-30 can confer 2-nonanone sensitivity to AWA and that the behavioral output depends on the identity of the neuron in which the receptor is expressed. Importantly, AWA-mediated attraction to diacetyl was not disrupted by ectopic *srh-30* expression (Fig. 4C), indicating that AWA function remained intact. Together, these results support the conclusion that SRH-30 acts as a sensory receptor for 2-nonanone and that neuron-specific expression of this receptor contributes to the aversive behavioral response.

**Figure 4.**
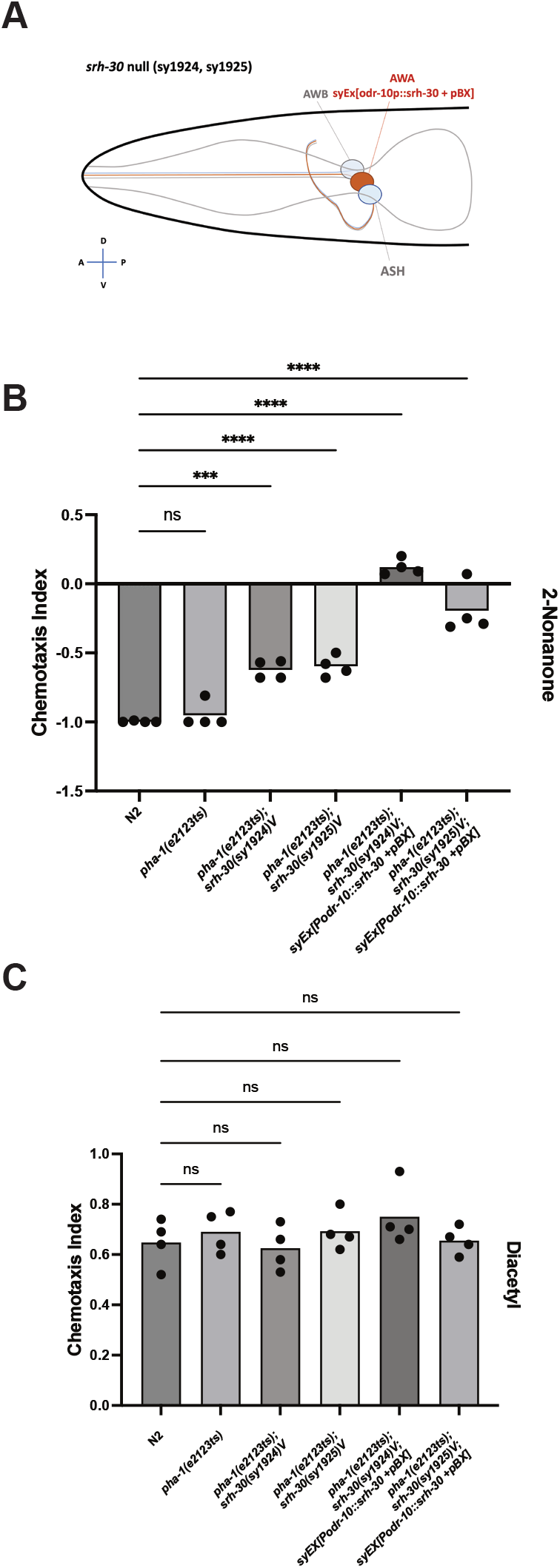
Ectopic expression of *srh-30* in AWA suppresses 2-nonanone avoidance in *srh-30* mutant animals. A) Schematic of ectopic *srh-30* expression in AWA using the AWA-specific *odr-10* promoter (Ex[*odr-10p*::*srh-30* + pBX]). B) Chemotaxis index to 2-nonanone. Ectopic AWA expression of *srh-30 (Ex[*odr-10p::srh-30 *+ pBX])* in *srh-30(sy1924)V* [PS9817] and *srh-30(sy1925)V* [PS9818] (two independent lines) shifts the aversive response toward neutrality. C) Chemotaxis index to diacetyl. Ectopic AWA expression of *srh-30 (Ex[*odr-10p::srh-30 *+ pBX])* does not alter attraction to diacetyl. Bars show mean ± SEM, dots indicate individual trials. Statistical significance was assessed by one-way ANOVA with Dunnett’s multiple-comparisons test relative to N2. Asterisks denote significance (* P < 0.05, ** P < 0.01, *** P < 0.001, **** P < 0.0001), ns indicates not significant.

## Discussion

Despite the large chemoreceptor repertoire of *C. elegans*, only a small number of odorant-receptor pairs have been experimentally defined[3, 12, 16]. Here, we identify SRH-30 as a chemoreceptor required for avoidance of 2-nonanone. By combining CeNGEN single-cell expression data with a focused reverse-genetic screen, we narrowed the large chemoreceptor repertoire to a tractable candidate list and identified *srh-30* as a gene whose loss reduces 2-nonanone avoidance, adding SRH-30 to the small set of experimentally matched odorant-receptor pairs in *C. elegans*.

Several lines of evidence support a sensory role for SRH-30. The original *srh-30* mutant showed reduced 2-nonanone avoidance but normal 1-octanol avoidance, arguing against a general chemotaxis defect. Two independent CRISPR-Cas9 null alleles reproduced the phenotype, arguing against background mutations in the original strain. A transcriptional reporter confirmed expression in AWB, the neuron previously implicated in 2-nonanone avoidance, and SRH-30::mNeonGreen localized to the distal tips of AWB sensory cilia, placing the protein in the appropriate compartment for odor detection. The mutant defect was partial, indicating that additional receptors in AWB, ASH, or other neurons may also contribute.

Ectopic expression of *srh-30* in AWA further supports this role. Because behavioral valence in *C. elegans* depends on the neuron in which a receptor is expressed, AWA-driven *srh-30* expression should shift the 2-nonanone response away from avoidance if SRH-30 confers ligand sensitivity. Consistent with this prediction, AWA expression of *srh-30* shifted 2-nonanone avoidance toward neutrality in *srh-30* null animals without affecting diacetyl attraction. The response was not converted to strong attraction, again reflecting possible parallel avoidance pathways of 2-nonanone receptors.

Together, these results identify SRH-30 as a receptor required for normal 2-nonanone avoidance and illustrate how single-cell transcriptomic atlases, combined with genetics and neuron-specific misexpression, can guide deorphanization of *C. elegans* chemoreceptors.

## Materials and Methods

### Strains and maintenance

*C. elegans* strains were maintained under standard conditions on nematode growth medium (NGM) agar plates seeded with *Escherichia coli* OP50. Unless otherwise indicated, animals were grown at 25°C to promote maintenance of injected extrachromosomal arrays in transgenic strains. Strains carrying the temperature-sensitive *pha-1(e2123)* allele, including *pha-1(e2123)*; *srh-30(sy1924) V* and *pha-1(e2123); srh-30(sy1925) V*, were maintained at 15°C. Bristol N2 was used as the wild-type reference strain. Candidate mutant strains were obtained from the Caenorhabditis Genetics Center (CGC), and strains used in the initial screen are listed in Supplementary Table 1. CRISPR-engineered strains were generated as described previously [26, 27]. Behavioral assays were performed using synchronized, age-matched animals 72 h after egg laying under 25°C.

### CeNGEN-based candidate selection

Single-cell RNA sequencing profiles for all 12 bilateral pairs of amphid neurons were obtained from CeNGEN (https://www.cengen.org; adult hermaphrodite dataset). To identify candidate chemoreceptors, genes expressed in both AWB and ASH were selected, while genes also expressed in other amphid neurons were excluded. Candidate genes with available mutant strains from the Caenorhabditis Genetics Center (CGC) were then prioritized for behavioral screening.

### Chemotaxis assay

Initial behavioral screening was performed using a quadrant avoidance assay on 10 cm assay plates. Each plate was divided into four quadrants. Odorant solution, 1 µL, was placed at positions A and C, and ethanol control, 1 µL, was placed at positions B and D. Animals were placed at the center of the plate and allowed to move freely for 20 min.

Assays were stopped by adding 200 µL chloroform to the plate lid. Animals remaining within the central 1 cm diameter circle were excluded from scoring.

The chemotaxis index was calculated as:

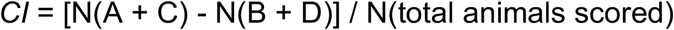

where N(A + C) represents the number of animals in the odorant quadrants and N(B + D) represents the number of animals in the ethanol control quadrants.

Validation assays for CRISPR-generated *srh-30* null mutants and transgenic animals were performed using a two-choice aversive chemotaxis assay on 10 cm assay plates. Plates were divided into odorant and control halves. Two 1 µL drops of odorant were placed on the odorant side, and two 1 µL drops of ethanol were placed on the control side. Animals were placed at the center of the plate and allowed to move freely for 20 min. Assays were stopped by adding 200 µL chloroform to the plate lid. Animals remaining within the central 1 cm diameter circle were excluded from analysis.

For the two-choice assay, the chemotaxis index was calculated as:

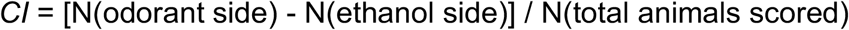

where N(odorant side) and N(ethanol side) represent the number of animals on the odorant and ethanol control sides, respectively.

For all assays, synchronized animals were generated by allowing day 1 adult hermaphrodites to lay eggs for 4 h at 25°C. Progeny were assayed 72 h after egg laying.

### CRISPR-Cas9 genome editing

Two independent *srh-30* null alleles, *srh-30(sy1924) V* and *srh-30(sy1925) V*, were generated using CRISPR-Cas9 genome editing. The Cas9 protein-based editing protocol was adapted from Paix et al. [26]. Guide RNAs targeting the *srh-30* coding region were designed using the *C. elegans* CRISPR guide RNA design tool [35]. Single-stranded donor oligonucleotides contained 35 bp homology arms flanking the edited region. WatCut (http://watcut.uwaterloo.ca/watcut/watcut/template.php?act=set_preferences), an online restriction analysis tool, was used to design restriction sites that did not alter the encoded protein sequence when applicable. crRNAs, tracrRNA, and donor oligonucleotides were commercially synthesized and dissolved in Nuclease-Free Duplex Buffer (Integrated DNA Technologies, Coralville, IA). Purified Cas9 protein was obtained from Prof. Tsui-Fen Chou. gRNA duplexes were generated by mixing crRNA and tracrRNA at a 1:1 ratio and incubating the mixture at 94°C for 2 min. Cas9 protein, 25 µM final concentration, and gRNA duplex, 27 µM final concentration, were mixed and incubated at room temperature for 5 min before donor oligonucleotides were added, 0.6 µM final concentration. Candidate edited animals were isolated as homozygous lines and verified by PCR and Sanger sequencing.

### Reporter and transgenic constructs

For expression analysis, approximately 3 kb upstream of the *srh-30* coding sequence was used to drive mKate2 expression, generating the *srh-30p::mKate2* reporter, syIs908. For protein localization, *srh-30* was fused to mNeonGreen under the control of the *srh-30* promoter, generating *srh-30p::srh-30::mNeonGreen*, syEx1951. For ectopic expression, *srh-30* was expressed in AWA using the *odr-10* promoter, generating *odr-10p::srh-30*.

### Transgenic animals

Standard *C. elegans* microinjection was used to generate transgenic animals carrying extrachromosomal arrays. Some arrays were subsequently integrated into the genome by X-ray irradiation [36].

### Fluorescence microscopy

Animals carrying fluorescent reporter constructs were mounted on 2% agarose pads and imaged using a Zeiss Axio Imager Z2 microscope equipped with an Apotome 2 and an AxioCam 506 mono camera. Images were captured using a Plan-Apochromat 63x/1.4 Oil DIC objective and ZEN Blue 2.3 software.

### Statistical analysis

Behavioral data are plotted as mean ± SEM, with individual dots representing independent trials where shown. Statistical comparisons were performed using one-way ANOVA with Dunnett’s multiple-comparisons test relative to N2, unless otherwise indicated. Significance levels are indicated as *P* < 0.05, *P* < 0.01, *P* < 0.001, and *P* < 0.0001. Exact sample sizes, *P* values, and the definition of an independent trial are provided in the corresponding figure legends.

## Acknowledgments

We thank the Caenorhabditis Genetics Center for providing strains. This work was supported by the National Institutes of Health Office of the Director (NIH/OD) under award number R24OD023041 and the National Institute of Neurological Disorders and Stroke (NIH/NINDS) under award number R01NS113119.

