## Supplemental Table 1 for "Identification of *srh-30* as a 2-nonanone receptor in *C. elegans*"

**Supplementary Table 1.**

| Gene | Gene ID | Strain |
|---|---|---|
| srh-18 | WBGene00005243 | VC20574; VC30144 |
| srh-30 | WBGene00005253 | VC20493 |
| Y40B10A.5 | WBGene00021490 | VC40563 |
| cyn-17 | WBGene00000893 | VC20057 |
| srw-139 | WBGene00005886 | VC40982 |
| C06A12.19 | WBGene00255702 | VC30012; VC20710 |
| srsx-33 | WBGene00044115 | VC40794 |

**Supplementary Table 1. CGC-available mutant strains for CeNGEN-derived candidate receptors express in AWB and ASH but not in other amphid sensory neurons.**
Candidate chemoreceptor genes were selected from CeNGEN single-cell RNA sequencing based on expression in the aversive amphid neurons AWB and or ASH, while excluding genes expressed in other amphid sensory neurons. For each candidate, the WormBase gene ID and the corresponding *C. elegans* Genetics Center (CGC) strain designation(s) used in this study are listed.
